# Assigning metabolic rate measurements to torpor and euthermy in heterothermic endotherms: “torpor”, a new package for R

**DOI:** 10.1101/717603

**Authors:** Nicolas J. Fasel, Colin Vullioud, Michel Genoud

## Abstract

Torpor is a state of controlled reduction of metabolic rate (*M*) in endotherms. Assigning measurements of *M* to torpor or euthermy can be challenging, especially when the difference between euthermic *M* and torpid *M* is small, in species defending a high minimal body temperature in torpor, in thermolabile species, and slightly below the thermoneutral zone (*TNZ*). Here, we propose a novel method for distinguishing torpor from euthermy. We use the variation in *M* measured during euthermic rest and torpor at varying ambient temperatures (*T_a_*) to objectively estimate the lower critical temperature (*T_lc_*) of the *TNZ* and to assign measurements to torpor, euthermic rest or rest within TNZ. In addition, this method allows the prediction of *M* during euthermic rest and torpor at varying *T_a_*, including resting *M* within the *TNZ*. The present method has shown highly satisfactory results using 28 published sets of metabolic data obtained by respirometry on 26 species of mammals. Ultimately, this novel method aims to facilitate analysis of respirometry data in heterothermic endotherms. Finally, the development of the associated R-package (torpor) will enable widespread use of the method amongst biologists.

**Summary statement:** The presented method and its associated R-package (torpor) enable the assignment of metabolic rate measurements to torpor or euthermy, ultimately improving the standardization of respirometry analyses in heterotherms.

## Introduction

Torpor is a state of controlled reduction of metabolic rate (*M*) and body temperature (*T_b_*) observed in numerous mammals and birds (Ruf and Geiser, 2015). This physiological state can occur over short periods (i.e. < 24h), often referred to as ‘daily torpor’, or it can last up to many days, often referred to as ‘hibernation’, which typically involves multiday torpor bouts separated by short spontaneous arousals (McKechnie and Lovegrove, 2002; Ruf and Geiser, 2015; Schleucher and Withers, 2001). The drop in *T_b_* may be considerable in hibernators, which often allow *T_b_* to reach values close to 0°C, but it can also be much shallower, as is often the case in species entering daily torpor (Ruf and Geiser, 2015). It has been suggested that a continuum may exist between these types of torpor as well as between torpor and euthermic rest (Boyles et al., 2013; McKechnie and Lovegrove, 2002; van Breukelen and Martin, 2015). Torpor use has profound implications on energy expenditure and allocation (Lyman et al., 1982) and affects many biological functions (Geiser and Brigham, 2012; Nowack et al., 2017). Accordingly, a vast amount of literature describes the occurrence of torpor and its associated energy savings in a multitude of species (Geiser and Ruf, 1995; Lovegrove, 2012; Nowack et al., 2020; Ruf and Geiser, 2015).

Determining whether measurements of *M* can be assigned to torpor or euthermy can however be challenging. The distinction between these two states is usually straightforward in typical hibernators, due to the more than 90% reduction in *M* common in these species when torpid at their usually hibernation ambient temperature (*T_a_*) (Geiser, 2004). Yet this distinction can become problematic when the difference between euthermic *M* (*M_e_*) and torpid *M* (*M_t_*) is small, which often occurs slightly below the thermoneutral zone (TNZ) (Geiser, 2011; Hainsworth and Wolf, 1970; Humphries et al., 2002; Speakman and Selman, 2003). Additional difficulties are encountered with species that either enter daily torpor, referred to as ‘daily heterotherms’, whose minimum *T_b_* in torpor often lies only slightly below the euthermic *T_b_* (Bartels et al., 1998; Bonaccorso and McNab, 1997; Genoud et al., 1990; McNab, 1980b), or who exhibit a large variability in *M* and *T_b_* at rest (Coburn and Geiser, 1998; Geiser et al., 1996).

Barclay, Lausen, & Hollis (2001) reviewed the variety of criteria that have been chosen in the past to distinguish torpor from euthermy. Apart from a minority of reports that identified torpor on the basis of behavioural features (Brice et al., 2002; Geiser and Kenagy, 1988; Geiser and Masters, 1994), the vast majority of studies have used patterns of variation in *T_b_* (or skin temperature) and/or *M* to separate the two states in the field or in the laboratory. Animals have been deemed to be in torpor below a threshold *T_b_* or *M* (Coburn and Geiser, 1998; Geiser et al., 1996; Hosken and Withers, 1999; Kelm and von Helversen, 2007; Levesque, 2008; Mzilikazi and Lovegrove, 2002), below some threshold temperature differential between the body and air (Levesque and Lovegrove, 2014), or below a threshold percentage of the euthermic rate of metabolism (Geiser, 1988a; Hudson and Scott, 1979). This diagnostic threshold value was sometimes calculated on the basis of the parameter’s variation in the euthermic state (e.g. Lovegrove & Raman, 1998; McKechnie, Ashdown, Christian, & Brigham, 2007), but has also been predicted by an equation based on body mass and *T_a_* (Willis, 2007). All available techniques have their limitations and are at least partly arbitrary (Barclay et al., 2001; Boyles et al., 2011), despite the efforts made to render them more objective.

With this paper, we propose an automated method for distinguishing torpor from euthermic rest in species that enter torpor, hereafter referred to as heterotherms following the terminology of Geiser & Ruf (1995) or Lovegrove (2012). Here, assignment to an either euthermic or torpid state is based on a probabilistic approach using the variation observed among measurements of *M* at varying *T_a_*. We assume that the relationship between *M* of resting animals and *T_a_* follows the classical “Scholander-Irving model” (Scholander, Hock, Walters, Johnson, & Irving, 1950; for a discussion see McNab, 2002), which was later extended to include torpor (Hainsworth & Wolf, 1970; Humphries, Thomas, & Speakman, 2002; Speakman & Thomas, 2003; Geiser, 2011). This model (Fig. 1) predicts the vast majority of patterns observed among endotherms entering daily torpor or hibernation. We explicitly consider that additional metabolic inhibition involving mechanisms other than the abolishment or reduction of thermogenesis necessary to maintain euthermy may occur during torpor (Geiser, 1988b; Geiser, 2004; Geiser and Kenagy, 1988; Guppy and Withers, 1999; Withers et al., 2016), by allowing the curve of torpor *M* to reach *T_lc_* at a level equal to – or lower than the resting metabolic rate within the *TNZ*. Thus, over all aforementioned techniques, our novel method provides major improvements, as it assigns measurements under an explicit framework (i.e. the extended Sholander-Irving model) and does so with a probabilistic approach based on observed variation.

**Figure 1:**
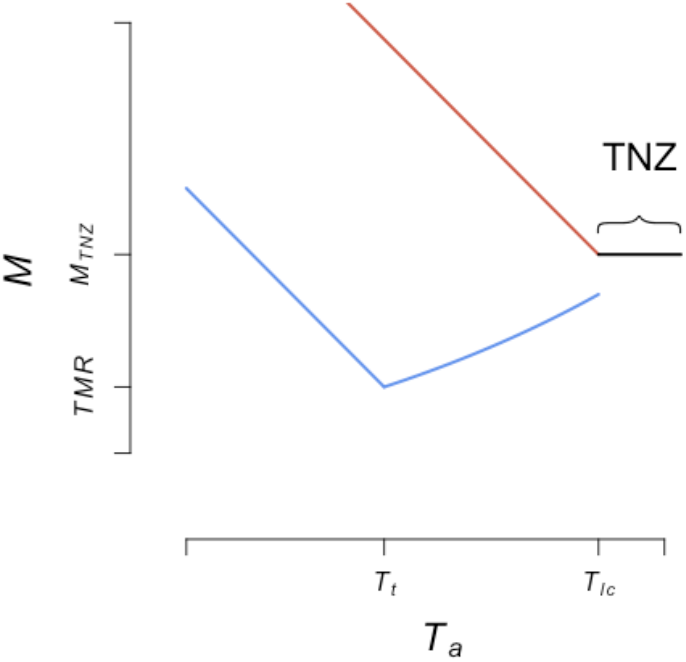
Representation of the relation between *M* during rest and torpor and *T_a_* on which the present method is based. Torpor is indicated in blue, euthermic rest below the *TNZ* in red and rest within the *TNZ* in black. The relation follows the classical Scholander-Irving model (Scholander, Hock, Walters, Johnson, & Irving, 1950; for a discussion see McNab, 2002), which was later extended to include torpor (Geiser, 2011; Hainsworth and Wolf, 1970; Humphries et al., 2002; Speakman and Thomas, 2003). We further consider the possible occurrence of additional metabolic inhibition (Geiser, 2004; Geiser and Kenagy, 1988; Withers et al., 2016), hence allow the torpor curve to reach *T_lc_* at a level equal to – or below *M_TNZ_*.

Our method is specifically intended to facilitate the discrimination between torpor and euthermy in laboratory experiments using respirometry. Additionally, it can also be useful to predict *M* at varying *T_a_*, by modelling the different parameters of the thermoregulatory curves. Further, we have assessed the performance of our method, by applying it to previously published data on the *M* of various mammals, including both heterotherms and species not undergoing torpor (hereafter referred to as homeotherms). Finally, we have provided a new package running within R (R Development Core Team, 2012) to allow researchers to apply our method to their own data. This package (“torpor”) comprises several useful functions that will improve standardization of the analyses of metabolic measurements for thermal biology.

## Results

Over all datasets, the proportion of assignments, that could be validly assigned (i.e. assignment confidence > 0.80, cf. “Assignment confidence” in section “Material and Methods”) ranged from 0.63 to 1.00 (median=0.87, Table 1). The corroboration index, which assesses the similarity between valid assignments made by our method and by the authors of the original datasets, ranged between 0.66 and 1.00 (median=1.00, Table 1). Complete matches between the method valid assignments and the authors’ descriptions were found for 15 out 28 datasets (e.g. *Nyctophilus geoffroyi* (Hosken and Withers, 1999), Fig. 2A). Mismatches mostly occurred close to *T_lc_* (e.g. *Peropteryx macrotis* (Genoud et al., 1990), Fig. 2C) but could also be found as far as 41.0°C below *T_lc_* (Fig. 3). Among homeotherms, no *M* measurement was assigned to torpor (e.g. *Sorex minutus* (Sparti & Genoud, 1989), Fig. 2B).

**Table 1:**
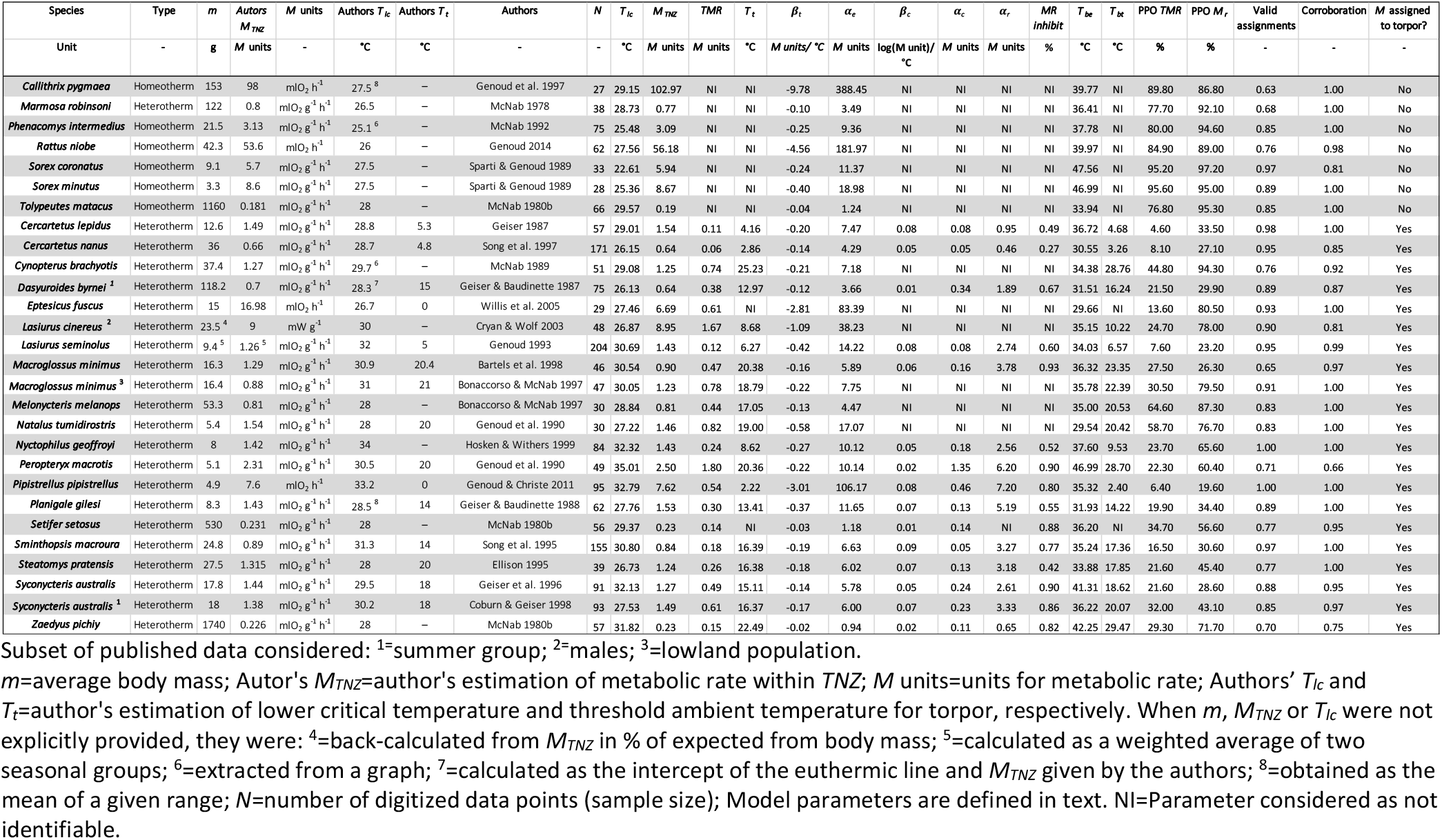
Model parameters and corroboration index for 28 datasets on small or medium-sized mammals described as either heterotherms or homeotherms.

**Figure 2:**
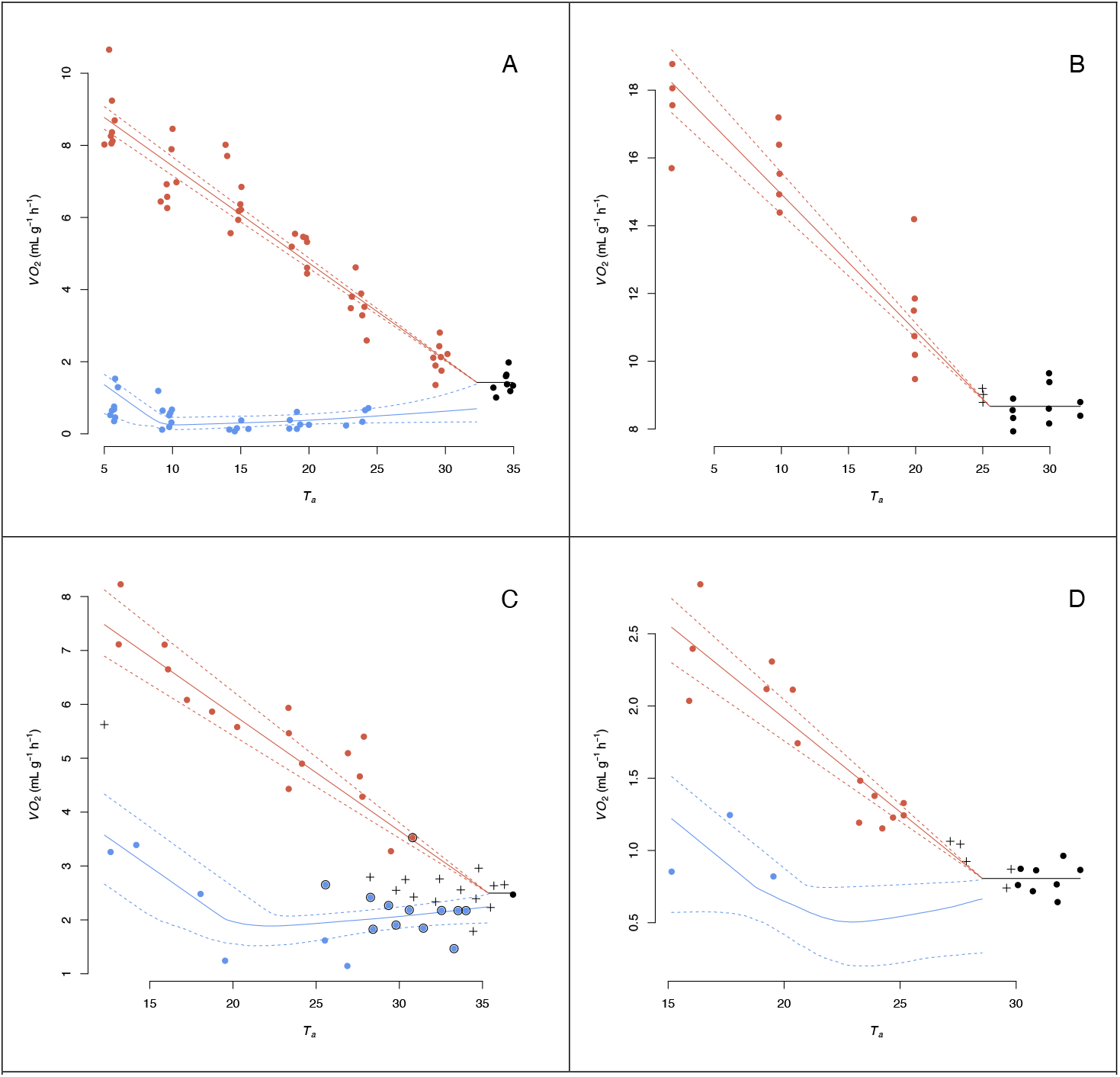
Four datasets of *M* measured at different ambient temperature (*T_a_*). Values were assigned to torpor (blue), euthermy (red) and *M_tnz_* (black) using the presented three-step method. Predicted values: median and 95% credible intervals are represented by continuous and segmented lines, respectively. Invalid assignments are highlighted with a cross and mismatches between authors and model assignments are surrounded with black circle. A/ Perfect corroboration between model and authors assignments in a heterotherm: *Nyctophilus geoffroyi* (Hosken and Withers, 1999). B/ Absence of mismatched assignment between model and authors assignments with some invalid assignments in a homeotherm: *Sorex minutus* (Sparti and Genoud, 1989). C/ Presence of mismatched or invalid assignments: *Peropteryx macrotis* (Genoud et al., 1990). D/ Absence of mismatched assignment between model and authors assignments with some invalid assignments, but insufficient number of torpor values to identify some torpor function parameters: *Melonycteris melanops* (Bonaccorso & McNab, 1997).

**Figure 3:**
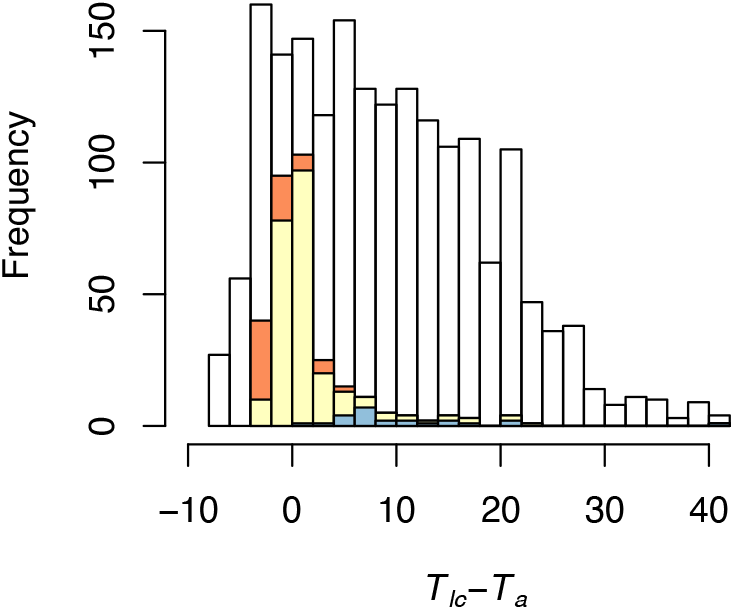
Frequency distribution of all the measurements from the 28 considered studies in relation to the difference between the method’s estimated *T_lc_* and the experimental *T_a_* (N=1898). Invalid assignments (i.e. assignment confidence < 0.8) are represented in yellow and assignments mismatches in orange and blue. Mismatches caused by the difference between the model’s estimated *T_lc_* and that of the authors are highlighted in orange and the remaining mismatches in blue.

In studies of homeotherms, where no *M* was originally assigned to torpor, prior and posterior distribution overlaps (*PPOs*) of *TMR* ranged from 76.80 to 95.60%. In identified heterotherms, *PPOs* of *TMR* values range from 4.60 to 64.60% (Table 1).

Modelled values of *T_lc_, M_TNZ_* and *T_t_* and extracted values from the original studies were significantly correlated (*T_lc_*: N=28, Pearson’s coefficient: 0.65, p<0.001, *M_TNZ_*: N=28, Pearson’s coefficient: 1.00, p<0.001, *T_t_*: N=14, Pearson’s coefficient: 0.97, p<0.001).

## Discussion

Our method aims to facilitate the discrimination between torpor and euthermy using the variation of *M* measured at varying *T_a_*. We tested it using 28 published sets of metabolic data obtained by respirometry on 26 species of small or medium-sized mammals. Selected species displayed a diversity of metabolic and thermal strategies ranging from permanent homeothermy to heterothermy including shallow, daily torpor and deep, long-term hibernation. The efficiency of our method proved satisfactory. Indeed, the corroboration index was generally high. Most conflicting assignments were a mere consequence of the difference between the estimated *T_lc_* and that defined by the authors (Table 1, Figs. 2C & 3). In particular, all mismatches between the model’s and the authors’ assignments concerning *M_e_* and *M_TNZ_* measurements in homeotherm species were explained by differences in estimated *T_lc_*. The remaining mismatches could generally be explained by intra-state frequency distributions of *M* that did not segregate clearly. This condition was often found close to *TNZ*, but also occurred in species where *T_t_* was relatively close to *T_lc_* (e.g. McNab, 1980a, 1989). Such discrepancies illustrate the difficulty of the assignment process when torpor and euthermy have to be distinguished solely on the basis of measurements of *M*. While the use of a statistical method enables an objective assignment, it is nevertheless worthwhile to recall that some authors also used patterns of *T_b_* to assign their data (e.g. McNab, 1980a, 1989). These specific cases should deserve further attention as they can highlight mechanisms decoupling *M* from the control of *T_b_* (e.g. Daniels, 1984; McNab, 1988; Heldmaier, Ortmann, & Elvert, 2004). One advantage of our method is to allow researchers to identify statistically data points that are difficult to assign. In this study, we considered assignments with a confidence lower than 0.8 as invalid. Obviously, an increase of this threshold value would lead to fewer assignment mismatches.

Our method also models the relationship between *M* and *T_a_* specific to each state, as well as several parameters describing the standard energetics of the animal(s) under study, including *M_TNZ_, TMR, T_lc_* and *T_t_*. Predicting *M* at any *T_a_* is crucial to model energy costs (e.g. Boyles, Johnson, Blomberg, & Lilley, 2020). It should however be recalled that our Bayesian inference-based method models all parameters whatever the available data. It remains therefore crucial to consider the parameters’ identifiability. The prior and posterior distribution overlaps (*PPO*) of *T_bt_* and *T_lc_* are not reliable for that purpose, because the prior distribution of those parameters largely depends on the data provided. From the *PPOs* of *TMR* obtained in studies where no *M* was assigned to torpor (i.e. “homeotherms”) and those obtained in identified heterotherms, we define a *PPO* higher than 75% as indicative of a parameter that is not identifiable.

The parameters estimated by our method correlated significantly with those extracted from the original studies. The number of *M_e_* values provided was probably not always sufficient to model *T_lc_* adequately. On one hand, a statistical estimation of *T_lc_* (e.g. in Song, Körtner, & Geiser, 1995, 1997; Bonaccorso & McNab, 1997; Genoud, Martin, & Glaser, 1997; Hosken & Withers, 1999; Willis, Lane, Liknes, Swanson, & Brigham, 2005; Genoud, 2014) was often lacking, leaving space for some potential misleading subjectivity by the authors of the original studies. On the other hand, an overestimation of *T_lc_* by our method remains possible if appreciable additional metabolic inhibition occurs during torpor (see below). The estimated *T_t_* was always close to that of the original studies. This higher precision in the estimation of *T_t_* in comparison to that of *T_lc_* is probably due to the lower variation of *M* exhibited during conforming torpor in comparison to that measured during euthermic rest. It is also not surprising that the *M_TNZ_* obtained by our method and by the authors of the original studies were close to identical, since both were calculated as the mean of nearly the same *M* values (i.e. all values above *T_lc_*)

An important aspect incorporated in our method is that we allow function for *M_t_* to be lower than *M_TNZ_* at *T_lc_*. We thus explicitly allow for additional metabolic inhibition to occur in some species during torpor, other than the cessation or reduction of the thermogenesis necessary to maintain euthermy (Geiser, 2004; Guppy and Withers, 1999; Withers et al., 2016). Caution should be kept in *M_r_* interpretation as an insufficient number of *M_t_* values corresponding to conforming torpor leads to the unidentifiability of this parameter (e.g. *Melonycteris melanops* (Bonaccorso & McNab, 1997), Fig. 2D). Consequently, the mentioned additional metabolic inhibition should only be considered when sufficient *M_t_* values corresponding to conforming torpor are provided and *M_r_ PPO* indicates adequate parameter’s identifiability. Considering only identifiable parameters (i.e. *Mr PPO* < 75%), we quantified additional metabolic inhibition as the fraction *M_r_* / *M_TNZ_*. Values ranged between 0.27 and 0.93 (N=18, median=0.77, Table 1). By extrapolation, we should consider the possibility that a hypometabolic state might occur within the *TNZ* as well (e.g. Grimpo, Legler, Heldmaier, & Exner, 2013; Reher, Ehlers, Rabarison, & Dausmann, 2018). With our method, we did not implement an assignment to such a hypometabolic state within the TNZ, hence *M_TNZ_* corresponds to the mean of all values from the *TNZ*. Our method might therefore underestimate the eumetabolic *M_TNZ_*, and might consequently overestimate *T_lc_*, if animals occasionally enter a hypometabolic state in the *TNZ,*. A predicted *M_t_* at *T_lc_* much lower than *M_TNZ_* should be viewed as an indication for a strong additional metabolic inhibition occurring during torpor and should invite for a careful consideration of the estimated *M_TNZ_* and *T_lc_*.

Throughout this paper, resting metabolic rate within the *TNZ* is referred to as *M_TNZ_* rather than to the basal rate of metabolism (*BMR*). Estimating the *BMR* requires several specific criteria: animals should be post-absorptive, adult, non-reproductive, and resting during a major inactive phase of the daily cycle (McNab, 1997). If these criteria are not met, the method still remains applicable, but the resting *M* estimated within the *TNZ* (i.e. *M_TNZ_*) will correspond to a minimal resting metabolic rate rather than to the *BMR*.

While the performance of our method appeared to be excellent with the selected datasets, its main limitations correspond to those of the model on which it is based. Indeed, although extremely fruitful, the Scholander-Irving model (Scholander et al., 1950) and its latter extensions (Geiser, 2011; Hainsworth and Wolf, 1970; Humphries et al., 2002; Speakman and Thomas, 2003) does not describe the thermal biology of all heterothermic species. Causes for significant divergence from the hereby model include a non-linear relationship between *M_e_* and *T_a_* (e.g. *Cercartetus nanus*: Song, Körtner, & Geiser, 1997), a difference in the slopes of the regressions of *M* versus *T_a_* in regulated torpor and euthermic rest (Geiser, 2004) and a strong dependence of *M_t_* on factors other than *T_a_*, such as body mass (Kelm and von Helversen, 2007) and duration since the last activity period or since last meal (Grigg et al., 1992; Morris et al., 1994). Furthermore, this model also does not describe adequately circadian (e.g. the shallow “rest-phase hypothermy” of many small birds (McKechnie and Lovegrove, 2002) or ultradian (Heldmaier et al., 1989) variations in *M*.

Our method is able to reveal the presence or the absence of distinct groups of values (euthermic rest vs torpor), but is not intended to judge whether any of the data points entered correspond to stable rates and/or to minimal values. Transitions between activity and rest or between euthermic rest and torpor as well as incomplete torpor bouts during which *M* never reaches a stable, minimal level (e.g. “test drops” (Hudson and Scott, 1979); see also (Genoud, 1993; Lyman, 1982)) should be excluded. It should also be recalled that long runs may be necessary for some heterothermic animals to achieve a state of steady torpor (e.g. Heldmaier, Ortmann, & Elvert, 2004). The above limitations should be kept in mind for a lucid application.

Then, individual variation is not explicitly considered in our method. Especially, body mass variation within a studied population might affect the assignment of *M* to torpor or euthermy in a complex manner (Genoud, 2014; Genoud et al., 2018; Glazier, 2005; Nespolo et al., 2003; Sassi and Novillo, 2015; Schleucher and Withers, 2001). However, there would be no particular difficulty to apply our method to single individuals, or repeatedly on single individuals.

Our method and the associated R-package (torpor) provide a way to standardize the analysis of respirometry data in relation to *T_a_*. Its major strength is that it uses a probabilistic approach to assign metabolic values to torpor or euthermy, rather than assigning them on the basis of a particular threshold value. Thus, the partly arbitrary nature of the assignment process is removed. Further, it can be applied to study intraspecific as well as interspecific variation in energetics. Parameters might now be extracted from the literature for comparative analysis, largely avoiding causes of variation due to the diversity of past assignment techniques. Ultimately, this method and the associated R-package (torpor) will ease intra- and inter-specific comparative analyses of endotherm energetics.

## Material and Methods

### The three-steps Method

Our present method is based on three steps and requires measurements of *M* of resting or torpid animals at *T_a_* ranging from below – to within the *TNZ*. Animals are assumed to be in one of three “states”: euthermic rest below the *TNZ* (*M_e_*), torpor below the *TNZ (M_t_*) or rest within the *TNZ* (*M_TNZ_*). We do not consider here the relation between *M* and *T_a_* above the *TNZ*. Consequently, metabolic rates measured at *T_a_* higher than the upper limit of the *TNZ* should be excluded. During the different steps, a mixture model based on Bayesian inference is run under varying conditions.

### The model

The Scholander-Irving model and its extensions (Fig. 1) consider that resting *M* measured within the *TNZ* is independent of *T_a_*. This rate is hereafter referred to as *M_TNZ_*. Below *T_lc_*, the *M* of euthermic animals (*M_e_*) increases linearly with decreasing *T_a_* :

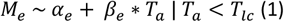

The model assumes that *M_e_* equals *M_TNZ_* at *T_lc_*, which enabled calculation of *α_e_*, the intercept of the line predicting *M_e_* at varying *T_a_*:

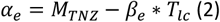

and of the slope of that line (*β_e_*):

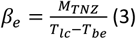

*T_be_* represents the hypothetical *T_a_* where *M_e_* equals 0, which correspond to the euthermic body temperature, provided that thermal conductance and body temperature do not vary with *T_a_* below the *TNZ* (McNab, 1980a).

Below the threshold *T_a_* separating regulated and conforming torpor (*T_t_*), the metabolic rate in torpor (*M_t_*) increases linearly with decreasing *T_a_* to defend the animal’s setpoint *T_b_* in torpor. That state, corresponding to *M_t_* measured at *T_a_* lower than *T_t_*, is usually referred to as “regulated torpor”:

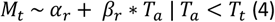

Where:

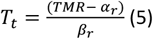

The intercept of the line for regulated torpor (*α_r_*) is obtained as:

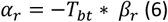

*TMR* represents the minimal *M_t_* measured at *T_t_*. *T_bt_* is the hypothetical *T_a_* at which *M_t_* of the regulated torpor would equal 0. *T_bt_* would correspond to the minimal body temperature in torpor (i.e. setpoint *T_b_* in torpor) provided that thermal conductance and body temperature during regulated torpor do not vary with *T_a_* below *T_t_*. Between *T_t_* and *T_lc_*, torpor is referred to as “conforming torpor” and *M_t_* follows an exponential curve:

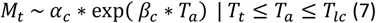

Where the coefficient of the exponent (*β_c_*) is calculated as:

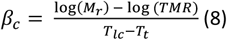

and the intercept for the exponential curve (*α_c_*) is calculated as:

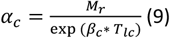

*M_r_* represents *M_t_* measured at *T_lc_*. In accordance with the general trend observed (Geiser, 2004), we assume that the slopes of the lines linking *T_a_* to *M_e_*(*β_e_*) and to *M_t_* of the regulated torpor, (*β_r_*) are similar:

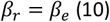

The following parameters are modelled: the fractions of measurements belonging to each state, which provides the value specific state membership probabilities for each measurement (see Second step: Metabolic rate (*M*) measurements pre-assignment, below), *T_be_, T_bt_, TMR*, and the standard deviations for *M_TNZ_* and regulated/conforming metabolic rates (*SD_TNZ_, SD_r_ and SD_c_*).

#### First step: Estimation of *M_TNZ_* and *T_lc_*

The method initially defines the highest possible *T_a_* within the dataset that still underestimates *T_lc_* (*T_lc_low_*). Above *T_lc_low_*, a linear regression between *M* and *T_a_* should neither result in a significant negative slope nor be affected by heteroscedasticity. Beginning with the ten *M* values measured at the highest *T_a_*, linear regressions are performed on sets of *M* values progressively including values at lower *T_a_*. The inclusion of additional *M* values obtained below *T_lc_* eventually leads to a significantly negative slope in the case of animals not entering torpor and/or to heteroscedasticity in the case of heterotherms. A significantly positive slope would reveal an absence or insufficient number of values for euthermic rest below the *TNZ*, hence it automatically stops the analysis as *T_lc_low_* is undefinable. Heteroscedasticity is assessed with a Breusch-Pagan test with the function “bptest” from package “lmtest” (Zeileis and Hothorn, 2002). Significance level for the Breusch-Pagan test is set at 0.05. The significance of the negative regression is assessed with a linear regression, function “lm” and significance level is set at 0.01.

Then, in order to get *T_lc_*, the model described in the second step (see Second step: Metabolic rate (M) measurements pre-assignment) is first run without data points measured above *T_lc_low_*. Moreover, within this step, the parameter *T_lc_* is also modelled. *M_TNZ_* provided in that analysis is the mean of the *M* measured at *T_a_* higher than *T_lc_low_*. Once *T_lc_* has been estimated, *M_TNZ_* is recalculated as the mean of the *M* measured at *T_a_* higher than *T_lc_*.

#### Second step: Metabolic rate (*M*) measurements pre-assignment

The assignment of *M* measurements to one of the three physiological states is based on a probabilistic process. Below *T_lc_*, a posterior categorical distribution is generated for each coupled data (*T_a_* and *M*), providing the probabilities to belong either to torpor or to euthermic rest (i.e. value specific state membership probabilities). Independent of the pre-assignment, *M* values higher than the predicted *M_e_* or lower than the predicted *M_t_* are automatically assigned to *M_e_* and *M_t_*, respectively. This latter automatic procedure is, however, only performed if at least one *M* measurement has been assigned to torpor during that second step. As defined previously, *M* values located above *T_lc_* are automatically assigned to *M_TNZ_*. Automatically assigned values get a state membership probability of one.

#### Third step: Metabolic rate (*M*) measurements final assignment and estimation of the functions parameters

In the third step, only *M* values situated between the predicted *M_e_* and *M_t_* are assigned. Measurements’ states that are automatically assigned during the second step are provided along with the coupled *M* and *T_a_* values. During that final step, the parameters of the functions relating *M* to *T_a_* below the *T_lc_*, as well as the standard deviation for the different physiological states are modelled.

### Bayesian parametrization

Prior distributions are defined either uninformatively or with biologically relevant limits. Except when specified, all prior distributions are Gaussian with a mean of 0 and a precision (i.e. 1/SD^2^) of 0.001 and are truncated based on the specified limits. Specifically, the prior distribution of *T_lc_* during the first step is constrained between *T_lc_low_* and the maximal *T_a_* recorded within the dataset. The prior distribution of *T_be_* is constrained between *T_lc_ (T_lc_low_* for the first step) and 50°C. The upper limit of the *T_bt_* prior distribution satisfies two conditions. First, *T_t_* should be inferior to the lower limit of the 95% credible interval (CI) of *T_lc_*. Second, the ratio of conforming *M_t_* values corresponding to body temperatures differing by 10°C (Q_10_) should not exceed a value of 5 (Geiser, 1988b). These conditions are verified by fixing the upper range of the prior distribution of *T_bt_* as:

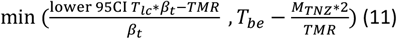

The lower limit of the *T_bt_* prior distribution is set at −5°C (Barnes, 1989). The upper limit of the prior distribution of *TMR* is defined as 80% of the *M_TNZ_* (Ruf and Geiser, 2015), while its lower one is fixed at 0. For *M_r_*, the prior distribution ranges from *TMR* to *M_TNZ_*. An uninformative Dirichlet distribution (i.e. two concentration values of 1) is used for the priors of the dataset specific state membership fractions, which provide the value specific state membership probabilities of each measurement (see Second step: Metabolic rate (*M*) measurements pre-assignment, above). Finally, the prior distributions of the standard deviations for *M_TNZ_* and regulated/conforming metabolic rates (*SD_TNZ_, SD_r_ and SD_c_*) are uniform, with that of *SD_r_* constrained between 0 and 3, that of *SD_c_* constrained between one fifth of the value of *SD_r_* and *SD_r_* and that of *SD_TNZ_* constrained between half the value of *SD_r_* and *SD_r_*.

Three different Markov chains are run during 50’000 iterations starting at initial values within the range of parameter space. The initial convergence phase is excluded by dropping the first 30’000 iterations. Markov chains are thinned by a factor of 10 and the Brooks–Gelman–Rubin criterion 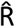 (Brooks and Gelman, 1998) is used to assess the convergence of chains (< 1.1). To ease parameters’ estimation, metabolic rate measurements are divided by their mean.

### Identifiability of the parameters

The prior and posterior distribution overlap (*PPO*) enables the evaluation of the relevance of some modelled parameters. Overlap values are obtained with the function “MCMCtrace” from package “MCMCvis” (Youngflesh, 2018). Truncated Gaussian prior distributions are provided with the function “rtruncnorm” from package “truncnorm” (Mersmann et al., 2018).

### Assignment confidence

The *M* value is assigned to the physiological state having the highest specific state membership probability. The assignment confidence represents the product of the highest specific state membership probability and of the probability that *T_a_* is above (for *M_TNZ_*) or below (for *M_t_* and *M_e_*) *T_lc_*. That latter probability is calculated from the variation in the estimation of *T_lc_* (cf. First step: Estimation of *M_TNZ_* and *T_lc_*), as the value at *T_a_* of the *T_lc_* cumulative posterior distribution function, or 1 minus that function respectively. A threshold proportion is then selected to consider any assignment with a significantly lower assignment confidence as invalid. That hypothesis is tested with a binomial test, function: “binom.test”, with the alternative hypothesis set as “greater”. The significance level for the binomial test is set at 0.05. For the present study, assignments with a confidence higher than 0.8 were considered as valid.

### Data collection and method evaluation

We evaluated our method by applying it to 28 published sets of *M* data obtained by respirometry. Mainly, we aimed at highlighting strengths and possible issues linked to the application of our method to typical sets of measurements. Thus, we were specifically interested in divergences between assignments made by our method and by the authors. The 26 species of small or medium-sized mammals investigated illustrate a diversity of metabolic and thermal strategies ranging from permanent homeothermy to heterothermy including shallow, daily torpor and deep, long-term hibernation. The selected datasets also differ in size (Table 1). For each published set, we considered all metabolic data provided, except those made above the upper critical temperature defined by the authors (see “the three-steps Method” above). In 20 species (here considered “heterotherms”, Table 1), rates of metabolism measured at *T_a_’s* below the described thermoneutral zone were originally assigned to either of two states, which were usually referred to as euthermy (or normothermy) and torpor. In *Cynopterus brachyotis*, values corresponding to particularly low *T_b_* were identified but not explicitly referred to as torpor (McNab, 1989). Some of the values obtained from torpid *Lasiurus seminolus* by Genoud (1993) were originally characterised by “irregular fluctuations” in *M* (Genoud, 1993). For simplicity, we treated these values as if the authors assigned them to torpor. The remaining six species are hereafter referred to as homeotherms (Table 1), as none of the values in the corresponding datasets were assigned to torpor in the original studies. Chosen studies reported *M_TNZ_* (mostly referred to as *BMR*) and *T_lc_* estimates, although the latter had to be extracted from a graph in two cases, or calculated as the intercept between the euthermic line below thermal neutrality and *M_TNZ_* provided by the authors in another case (Table 1). In all dataset the authors provided a sufficiently precise graph of metabolic rate values as a function of *T_a_* and several of them also estimated *T_t_*. Plotted data were digitized using the software Plot Digitizer (Huwaldt and Steinhorst, 2015) except for the *Pipistrellus pipistrellus* dataset, which was provided by MG. Sample sizes of the digitized data ranged between 27 and 204 (Table 1). We tested the correlation between the *M_TNZ_, T_lc_* and *T_t_* values modelled with the present method and those provided by the authors with Pearson’s paired-samples correlation test, function “cor.test”. Cases in which the estimation of *T_t_* fell outside the range of measured *T_a_* were excluded from this analysis as *T_t_* was considered unidentifiable. Then, we examined whether the assignment of the metabolic values to torpor or euthermy made by the authors and provided by the method coincided, and we calculated a corroboration index as the fraction of the matched assignments. Only measures with valid assignments (i.e. assignment confidence > 0.80, cf. “Assignment confidence” in previous section) were considered for the correlation tests and the corroboration index calculations. Finally, in order to define a *PPO* range highlighting an identifiable parameter, the *PPOs* of *TMR* from studies where part of the *M* measurements were assigned to torpor were compared with those from studies where no *M* measurement was assigned to torpor. In those later studies *TMR* was considered unidentifiable.

## Supporting information

SuppInfo_exemple

## List of main abbreviations

*BMR*: Basal metabolic rate.
*M*: Resting rate of metabolism, expressed in W, mlCO_2_ h^−1^ or mlO_2_ h^−1^. Often expressed as a mass-specific rate (e.g. in W g^−1^).
*M_TNZ_*: Resting rate of metabolism within the *TNZ*.
*M_e_*: Euthermic *M*.
*M_r_*: predicted *M_t_* at *T_lc_*. *M_r_* may be lower than *M_TNZ_* if additional metabolic inhibition is involved, other than the abolishment or reduction of thermogenesis necessary to maintain euthermy.
*M_t_*: Torpid *M*.
*PPO*: prior and posterior distribution overlap.
*T_a_*: Ambient temperature.
*T_b_*: Body temperature.
*T_lc_*: lower critical *T_a_* or lower limit of the *TNZ*.
*TMR*: minimal metabolic rate in torpor, measured at *T_t_*.
*TNZ*: Themoneutral zone. The range of *T_a_* where *M* is independent of *T_a_*.
*T_t_*: Threshold *T_a_* separating regulated from conforming torpor.

## Acknowledgements

We are thankful to Philippe Christe and Nicolas Salamin for their support during the elaboration of the model. We also thank four anonymous reviewers who contributed to improve the original manuscript.

## Competing interests

No competing interests declared

## Funding

This study was financially supported by the Department of Ecology and Evolution of the University of Lausanne (Switzerland) and by the Swiss National Science Foundation (grant number: P2BEP3_168709 to NJF)

## Notes

### Competing Interest Statement

The authors have declared no competing interest.

