## Supplementary material for "Assigning metabolic rate measurements to torpor and euthermy in heterothermic endotherms: “torpor”, a new package for R": SuppInfo_exemple

### An example

#### How to use torpor ? An example

This vignette presents a suggested workflow of the `torpor` package. This package is firstly aimed at users not familiar with `rjags`, yet interested in investigating the thermoregulatory pattern of their model species, especially when the latter is a heterotherm and when metabolic rate measurements are to be assigned to either torpor or euthermy.

This workflow is based on one of the provided datasets. We go through a complete analysis to illustrate how to use the package functions. At each step, we also present some options for the users and the possible solution to expected problems. Note, however, that this example does not present all aspects of the package.

##### Installing the package

As the package is still not available on CRAN, users have to download it by calling the following command.

```
remotes::install_github("vullioud/torpor", build_vignettes =
T,force=T)
```

The program JAGS can be downloaded from the website: https://mcmc-jags.sourceforge.io/

##### Fitting a model

The first step in any analysis is to import the data. In order to fit a model using the function `tor_fit()`, we need a set of measurements of Metabolic rate $M$ and ambient temperature $T_a$ at which the measurements have been recorded. These values can be vectors of the same length or two columns of a data frame. In the chosen example on the Tasmanian pygmy possum, *Cercartetus lepidus*, we have 103 metabolic rate values at various $T_a$ obtained by digitalization of a figure (Geiser, 1987).

The data are accessible with the following command.

```
library(torpor)
data(test_data2)
str(test_data2)
```

```
## 'data.frame':    103 obs. of  2 variables:
##  $ Ta    : num  2.82 3.01 3.09 3.31 3.33 ...
##  $ VO2ms: num  0.364 NA NA NA 0.222 ...
```

The lower critical temperature ($T_{lc}$) of the thermoneutral zone (TNZ) and the metabolic rate within TNZ ($M_{TNZ}$) can be estimated by or provided to the `tor_fit()` function. These two values were originally estimated by the author and can be found in the documentation of the dataset (test_data2). For the present example, we will estimate $T_{lc}$ and provide $M_{TNZ}$. `tor_fit()` represents the core function of the model and should be the first step in any analysis using the torpor package.

The model is fitted with the following call:

```
model <- tor_fit(Ta = test_data2$Ta,
                 M = test_data2$VO2ms,
                 Mtnz = 1.8,
                 confidence = 0.5)
```

The output of the `tor_fit()` function is a list containing information on the posterior distributions of the investigated parameters as well as on the convergence of the chains and on the prior-posterior distributions overlap of some parameters. These information can also be called by the function `tor_summarise()`. The rest of the package offers several functions to make more sense of the model output, to check its consistency and to represent it graphically. Researchers familiar with `jagsUI` can develop their analysis from that point.

#### Checking the adequacy of the model

Once the model is fitted, it is recommended to verify the adequacy of the results. This can be achieved via the function `tor_summarise()`. The latter will return a list with the essentials: A dataframe with the mean and median of parameter estimates, the 95% credible interval and the $\hat{R}$ value (i.e. chain convergence estimation). It also reports the parameters' identifiability.

```
summary <- tor_summarise(model)
```

The parameter estimates are accessible with:

```
summary$params

   parameter   mean CI_2.5 median CI_97.5  Rhat
1       tau1  0.139  0.094  0.132   0.219 1.006
2       tau2  0.424  0.333  0.420   0.545 1.002
3       tau3  0.132  0.090  0.129   0.193 1.002
4       inte  7.396  7.090  7.396   7.692 1.000
5       intc  0.092  0.073  0.087   0.134 1.002
6       intr  0.823 -0.773  0.962   1.272 1.005
7      betat -0.197 -0.207 -0.197  -0.186 1.000
8      betac  0.069  0.036  0.071   0.093 1.001
9         Tt  3.586 -4.469  4.297   5.828 1.005
10       TMR  0.118  0.092  0.116   0.148 1.002
11        Mr  0.667  0.322  0.663   1.092 1.002
12       Tbe 37.587 37.121 37.580  38.110 1.000
13       Tbt  4.186 -3.939  4.900   6.455 1.005
14       Tlc 28.571 26.421 28.434  31.608 1.019
15      Mtnz  1.800     NA  1.800      NA    NA
```

#### Checking the convergence

If the $\hat{R}$ values given in the summary dataframe are larger than 1.1, we recommend to refit the model and increase the number of iterations and burn-ins. This can be done by setting the parameter `fitting_options = list(ni = , nb = )`. It is also possible to look graphicaly at the convergence using the `jagsUI` function `traceplot()`. Let's have a look at the convergence for the parameters `betat` and `betac`.

```
jagsUI::traceplot(model$mod_parameter, c("betat", "betac"))
```

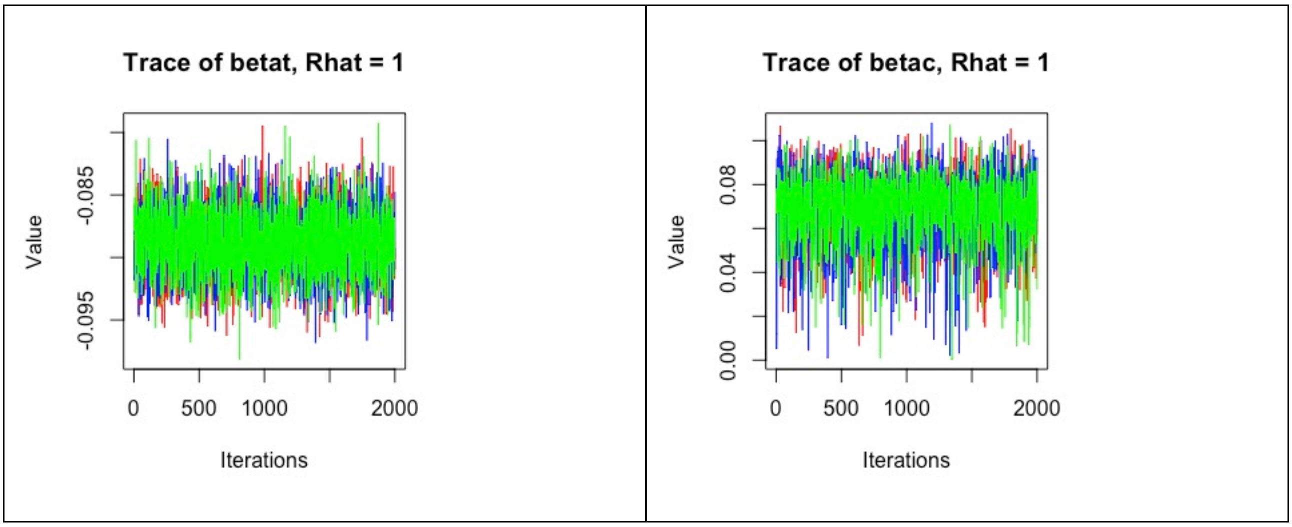


#### Checking the parameters identifiability

In addition to a verification of the convergence it is advised to control the identifiability of some parameters. This is done by comparing the prior and the posterior distributions. A warning will be sent to the user for estimated parameters whose PPO are higher than 75% (Fasel et al. in prep.). Parameter identifiability is provided for $T_{lc}$, $M_r$, and *Tbe*, *TMR* and *Tbt*. The ovelap is also given by the `tor_summarise()` function. Let's continue with our analysis of *Cercartetus lepidus*:

```
summary$ppo

  name  ppo
1   Mr 31.2
2  TMR  4.6
```

#### Making sense of the model

Once the basic checks have been done, we can go on with the evaluation of the output. There are two main functions that deal with predictions. `tor_assign()` firstly gives the classification of the raw data based on the predictions of the model. `tor_predict()` further gives the predicted $M$ in torpor and euthermy for a given $T_a$. Finally, we can plot the result using the function `tor_plot()`.

#### Getting the assignments

To look at the assignments of the data and the corresponding predicted values we use the function `tor_assign()`, which will assign the measured metabolic values to either torpor or euthermy and returns a dataframe with the measured $M$, the measured $T_a$, the predicted $M$ and the assignment. (predicted_state).

```
classification <- tor_assign(model)

head(classification)

  measured_M measured_Ta assignment predicted_M
1  0.3643880     2.81631     Torpor   0.4094219
2  0.2222220     3.32812     Torpor   0.3084751
3  0.5080130     3.37791     Torpor   0.2985989
4  0.4407550     3.94446     Torpor   0.1889358
5  7.0744500     4.55155   Euthermy   6.5006503
```

```
6  0.0538837     4.60326     Torpor   0.1252308
```

#### Prediction

The `tor_predict()` function is slightly different as it takes a vector of $T_a$ as input and returns the predicted *M* both in torpor and in euthermy and the 95% credible interval. For example, let's see what are the predicted *MR* at Ta 22°C.

```
prediction <- tor_predict(model, 22)

head(prediction)

Ta assignment      pred    upr_95    lwr_95
1 22     Torpor 0.4200096 0.6094252 0.2516173
2 22   Euthermy 3.0664153 3.1332578 2.9970233
```

#### Plotting the data and predicted values

Finally a built-in function allows plot the results. The user can modulate labels (with `xlab` and `ylab`), colors (with `col_torp`, `col_eut` and `col_Mtnz`) and can save the plot using the arguments `pdf = TRUE`.

```
tor_plot(
  tor_obj = model,
  col_torp = "cornflowerblue",
  col_eut = "coral3",
  ylab = "M",
  xlab = "Ta")

title(main="Cercartetus lepidus")
```
